# Reciprocal cross-feeding between bacteria can limit the emergence of metabolic dependencies

**DOI:** 10.1101/2024.08.19.608702

**Authors:** Ying-Chih Chuang, Megan G. Behringer, Gillian Patton, Jordan T. Bird, Jeffrey L. Mazny, Jennifer R. Gliessman, Crystal E. Love, Ankur Dalia, James B. McKinlay

**Author notes:** Corresponding author: James B. McKinlay: **Email:**. Current address: Department of Biological Sciences, University of Southern California, Los Angeles, CA, USA.

## Abstract

Cross-feeding is prevalent in microbial communities. Through time, cross-feeding is thought to enrich for loss-of-function mutations, thereby creating or reinforcing dependencies between community members. However, few studies have compared how cross-feeding affects the evolutionary trajectory of partners compared to monoculture conditions. Here we compared mutations that were differentially enriched in bacterial monocultures versus cocultures pairing phototrophic *Rhodopseudomonas palustris* with fermentative *Escherichia coli* in an obligate cross-feeding relationship based on the exchange of nitrogen and carbon for 650-800 generations. Opposite trends for the number of differentially enriched mutations were observed for each species; *R. palustris* accumulated more unique mutations in monoculture whereas *E. coli* accumulated more unique mutations in coculture. Contrary to expectations, the emergence of additional dependencies was more apparent for both species in monoculture, even though additional layers of cross-feeding involving iron and adenine were present in coculture. We reasoned that iron and adenine cross-feeding occurred at levels sufficient to repress gene expression in the recipient, thereby promoting gene retention by lowering gene cost. Thus, the influence of cross-feeding on evolutionary trajectories can vary with organisms and conditions, and there are situations where cross-feeding can limit, rather than promote, emergent dependencies.

**IMPORTANCE:** Bacteria commonly engage in cross-feeding where nutrients are transferred between neighbors. Cross-feeding is thought to alleviate energy expenditures for genes whose role can be met by cross-fed nutrients, leading to eventual gene loss. However, few examples have been documented, especially in comparison to monocultures where bacteria are grown without a cross-feeding partner. Here we grew cocultures pairing phototrophic *Rhodopseudomonas palustris* with fermentative *Escherichia coli* alongside corresponding monocultures for 650-800 generations. While coculture conditions required obligate exchange of nitrogen and carbon, additional cross-feeding of adenine and iron likely occurred. Contrary to expectations, iron and uncharacterized dependencies emerged in monocultures but not in cocultures. Low expression for iron scavenging and adenine synthesis genes in cocultures suggested that cross-feeding repressed gene expression, thereby lowering gene cost. Thus, whereas there are likely cases where cross-feeding makes costly genes dispensable, there are also cases where cross-feeding lowers gene cost, thereby promoting gene retention.

## INTRODUCTION

Microbial metabolism shapes local chemical environments in ways that elicit regulated responses from neighboring cells, or even select for adaptive mutations. In some cases, extracellular metabolites can be used as nutrients by neighboring cells, establishing cross-feeding (1). Cross-feeding can lead to obligate dependencies through adaptive loss-of-function (LOF) mutations, when the benefit of exploiting an extracellular metabolite outweighs the cost of retaining a biosynthetic gene (2). When the recipient benefits at the expense of the producer, the recipient can be labeled a ‘cheater’ but when the recipient does not detrimentally affect the producer, the recipient can be labeled a ‘beneficiary’ (2, 3). The latter is described in the Black Queen Hypothesis (BQH), wherein cross-feeding provides a cost-benefit advantage that promotes adaptive gene-loss (2). The BQH is commonly used to explain the prevalence of auxotrophs, which cannot synthesize one or more essential nutrients (4, 5). Indeed, several groups observed that auxotrophs with biosynthetic gene deletions have a fitness advantage over their prototrophic counterparts when corresponding biosynthetic precursors were provided (6–10).

Whereas cross-feeding can promote adaptive gene loss, gene loss is also potentially constrained during cross-feeding by several factors. For example, the energetic costs of N_2_ fixation, converting N_2_ into 2 NH_4_^+^ via nitrogenase, was predicted to be under pressure for gene loss because NH_4_^+^ can escape and potentially enable higher growth rate for recipients (2). However, we found that growth rates supported by leaked NH_4_^+^ did not exceed those possible via N_2_-fixation due to low NH_4_^+^ availability relative to N_2_, even when bacteria were engineered to excrete NH_4_^+^ (11). Moreover, as has been documented with cheaters (12–14), nutrient availability can repress gene expression and thereby lower gene cost (4, 13, 15); most of a gene’s cost comes from its expression (16–20). Thus, repressing gene expression could have similar cost-savings as a LOF mutation and thereby decrease selective pressure for LOF mutants.

Previously we engineered cocultures of phototrophic *R. palustris* and fermentative *E. coli* to study the mechanisms and evolution of obligate cross-feeding (21–25). Using anoxic conditions, *E. coli* ferments glucose into organic acids that provide essential carbon to *R. palustris* while *R. palustris* converts N_2_ into NH_4_^+^, providing essential nitrogen to *E. coli* (21). The two species are not believed to interact in nature, providing an opportunity to observe how the relationship evolves without pre-adaptation (22).

Whereas the coculture was engineered to rely on obligate NH_4_^+^ cross-feeding, other interaction layers exist. For example, we discovered that *R. palustris* externalizes enough adenine to support an *E. coli* purine auxotroph population (26, 27). How such secondary interactions affect the evolutionary trajectory of each species is unknown.

Here we compared differential mutation accumulation in *E. coli* and *R. palustris* monocultures versus cocultures for >600 generations. The number of differentially enriched mutations in monocultures versus cocultures was opposite for each organism. For several mutations, we performed a preliminary investigation to gain insights into why they were differentially enriched. In some cases, we correlated a lack of dependency emergence in coculture to inhibitory effects of cross-feeding on gene expression. Our results offer diverse insights into how cross-feeding influences evolutionary trajectories and caution against general expectations for dependencies through LOF mutations.

## RESULTS AND DISCUSSION

### *R. palustris* and *E. coli* can coexist for hundreds of generations

We established long-term, batch, anaerobic (i) monocultures of phototrophic, N_2_-fixing, NH_4_^+^-excreting *R. palustris* NifA* (CGA676) (28), supplied with a mixture of organic acid salts (neutral pH), ethanol, and bicarbonate to mimic *E. coli* fermentation products, (ii) monocultures of fermentative *E. coli* PFM2, a MG1655 derivative with repaired *rpoS* and *rph* genes (29, 30), supplied with 10 mM glucose and 10 mM NH_4_Cl, and (iii) cocultures supplied with 25 mM glucose (Fig 1A). All cultures were constantly illuminated, and 2.5% v/v was transferred every seven days. All conditions included an exponential growth phase but only monocultures experienced a nutrient-limited stationary phase. In cocultures, glucose and consumable organic acids remained at the time of transfer and likely supported growth, however slow due to the low pH. Ultimately, all conditions were serially transferred for 650 – 803 generations (∼5 generations per transfer; Fig 1).

**Fig 1.**
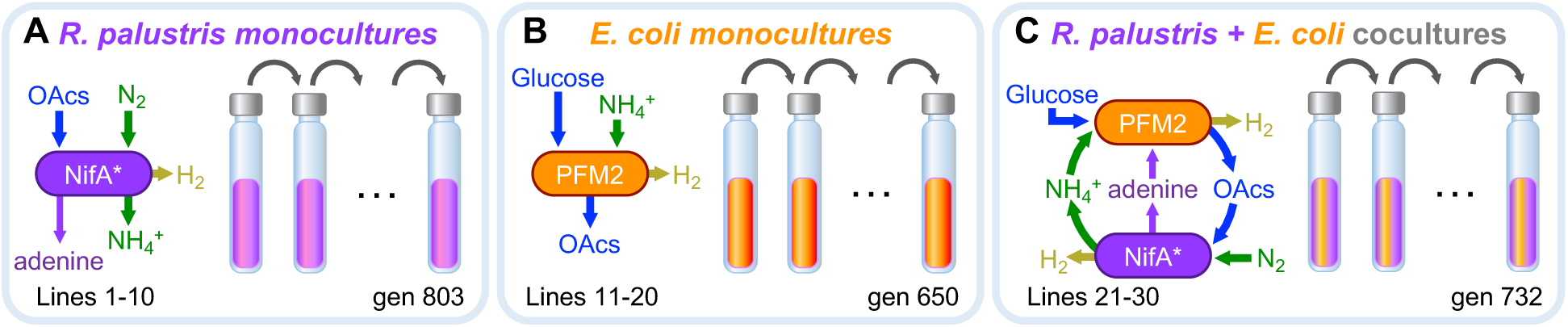
Long-term anaerobic monoculture and coculture conditions. All cultures were laid flat without shaking and were constantly illuminated. **(A)** Phototrophic *R. palustris* NifA* (CGA676) was fed a mixture of organic acids (OAcs), ethanol, and NaHCO_3_ to mimic carbon sources available from *E. coli* fermentation products. The NifA* mutation causes NH_4_^+^ excretion during N_2_ fixation. H_2_ is also produced as a co-product of N_2_ fixation. *R. palustris* also excretes adenine independent of the NifA* mutation. **(B)** *E. coli* PFM2 was fed 10 mM NH_4_Cl and 10 mM glucose, which it fermented to a mixture of OAcs, ethanol, CO_2_, and H_2_. **(C)** Cocultures were fed 25 mM glucose. Growth of each species is dependent on the exchange of NH_4_^+^ and OAcs.

We sequenced gDNA from populations at several timepoints. We then compared mutations that were differentially enriched in monoculture versus coculture by applying following criteria: (i) exclude synonymous mutations, (ii) no exact mutation in the opposing monoculture or coculture condition, (iii) observed in the last two sequenced timepoints, (iv) with a frequency >0.1 in the last timepoint. These criteria resulted in a list of 167 mutations from *R. palustris* monocultures, 31 from *E. coli* monocultures, 43 from *R. palustris* in cocultures, and 132 from *E. coli* in cocultures (Fig 2A, Table S1-5). These counts include redundancy, where some mutations were observed in multiple lines. In all conditions, most mutations were nonsynonymous, except for *E. coli* monocultures, where most mutations were small insertions/deletions (Fig 2B). Mutations due to mobile genetic elements were only observed in *E. coli* (Fig 2B). To narrow our focus on positive selection, we added a fifth criteria of parallelism where the same gene or intergenic region must be mutated in at least two lineages.

**Fig 2.**
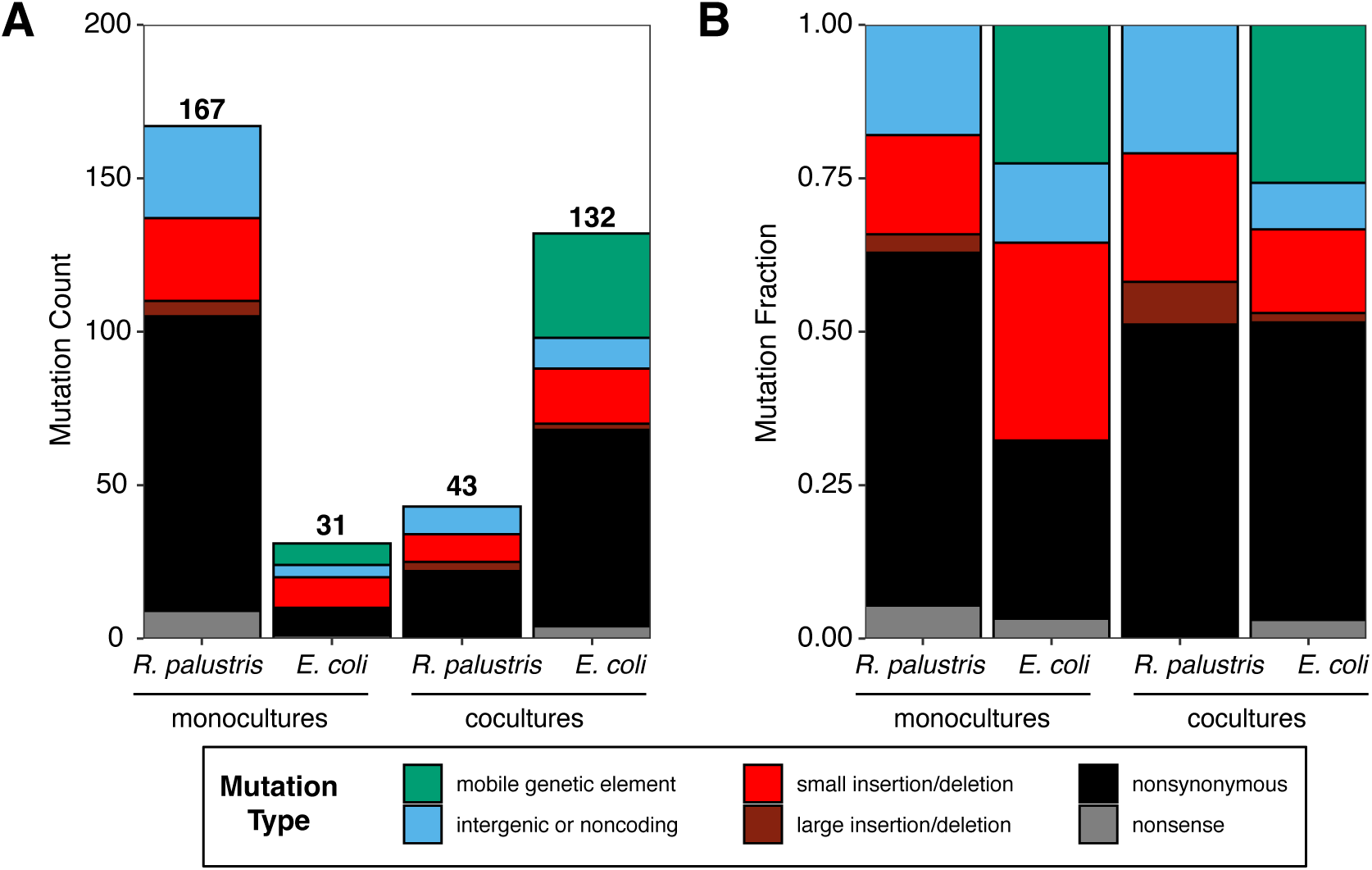
Differentially enriched mutations in long-term monocultures and cocultures. Mutation counts and fractions are the sum of mutations across all 10 lines in each condition. To be counted, mutations had to meet the following criteria: (i) exclude synonymous mutations, (ii) no exact mutation in the opposing monoculture or coculture condition, (iii) observed in the last two sequenced timepoints, (*Rp* monoculture, generations 781 and 803; *Ec* monoculture, generations 500 and 650; coculture, generations 655 and 732), (iv) with a frequency >0.1 in the last timepoint.

### Differentially enriched *R. palustris* mutations mostly occurred in monocultures

Most differentially mutated *R. palustris* genes and intergenic regions occurred in monocultures: 19 in monoculture vs 5 in coculture (Fig 3). We considered that the higher mutation frequency in monoculture could be due to a *mutS,* encoding a mismatch repair protein; lines 2 and 7 had a *mutS* mutations that met our criteria and most other lines had a low frequency *mutS* mutation at some point during the experiment.

**Fig 3.**
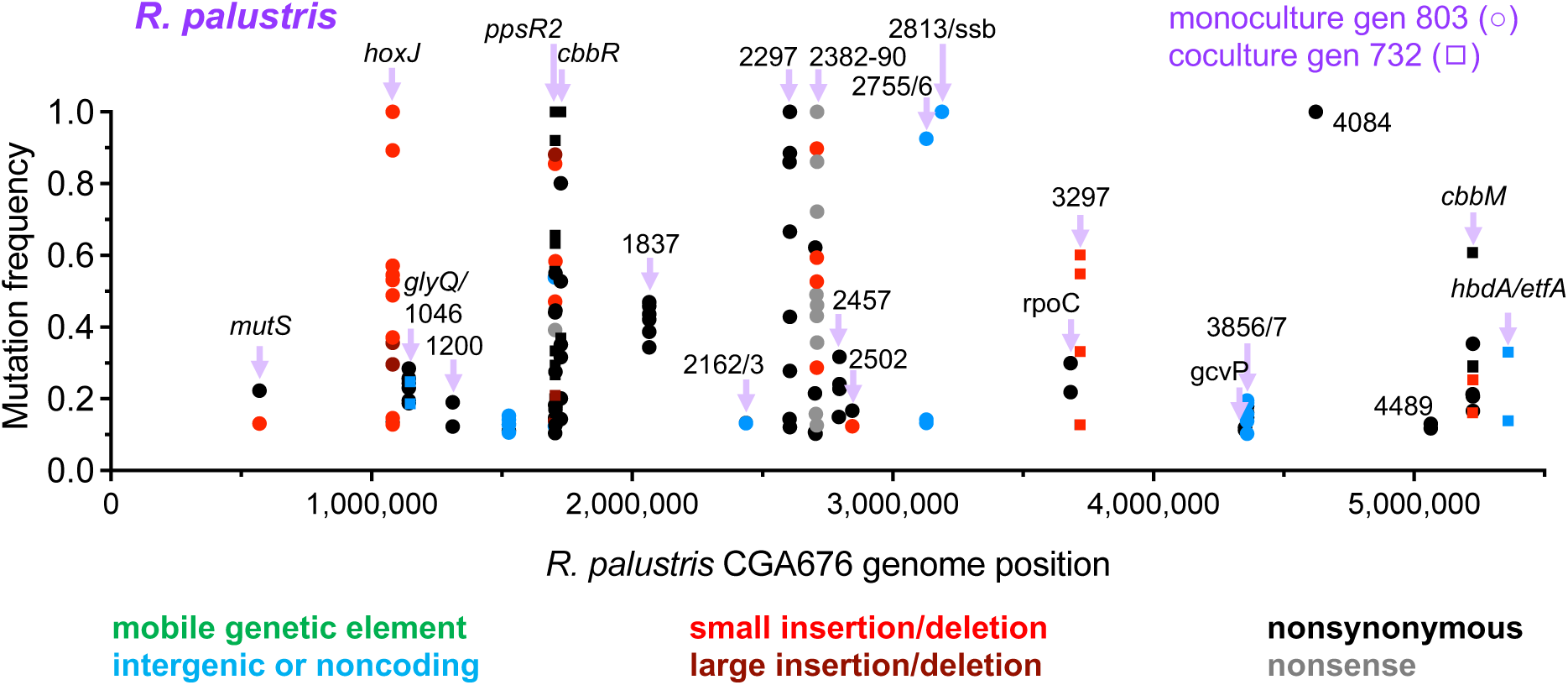
Mutations enriched in *R. palustris* NifA* in long-term monocultures and cocultures. Each point represents a mutation in a single lineage. Mutations are shown if they are not a synonymous mutation and were observed: (i) exclude synonymous mutations, (ii) no exact mutation in the opposing monoculture or coculture condition, (iii) observed in the last two sequenced timepoints, (*Rp* monoculture, gen 781 and 803; *Rp* coculture, gen 655 and 732), (iv) with a frequency >0.1 in the last timepoint, and (v) the same gene or intergenic region was mutated in at least two lines for a given condition. Generation: gen.

However, the total number of differentially-enriched mutations was similar across *R. palustris* monoculture lines (range: 8-23; Table S1) with only 6 of 21 mutations in line 2 and 11 of 23 mutations in line 7 potentially attributed to *mutS*, since they occurred at a lower frequency than the *mutS* mutation (*mutS* mutation frequency was 13 and 22% in lines 2 and 7, respectively; Table S1). Thus, *mutS* mutations did not likely drive many of the other mutations.

Several differentially mutated genes in coculture also acquired different mutations in monoculture. Among these was *ppsR2*, encoding an O_2_-responsive regulator of genes for phototrophic growth. *ppsR2* is commonly mutated in *R. palustris* because they provide an advantage under low-light conditions, which occurs at high cell densities (31). Other genes that were commonly mutated in monoculture and coculture were *cbbM*, encoding the Calvin cycle type II Rubisco and *cbbR*, encoding a Calvin cycle regulator. It is unclear why mutations were enriched in these genes, but it is possible that they are associated with the role of CO_2_ fixation in electron balance during photoheterotrophic growth or electron distribution between CO_2_ and N_2_ fixation (28, 32). We did not investigate Calvin cycle gene mutations further herein.

### *R. palustris* monocultures evolved to utilize H_2_

Both species produce H_2_, *E. coli* via fermentation and *R. palustris* as a coproduct of nitrogenase activity (33). Curiously, H_2_ levels decreased through serial transfers in *R. palustris* monocultures but not in cocultures (Fig 4A, B). A commonly mutated gene in *R. palustris* monocultures was *hoxJ,* which encodes a sensor kinase that governs H_2_-oxidizing hydrogenase by repressing the transcriptional activator HoxA (Fig 4C). Normally, HoxJ repression of HoxA is relieved when HupUV senses H_2_. However, the *R. palustris* ancestor had a *hupV* mutation, causing HoxJ to constitutively repress HoxA (34). Deleting HoxJ was previously shown to lead to hydrogenase activity (34).

**Fig 4.**
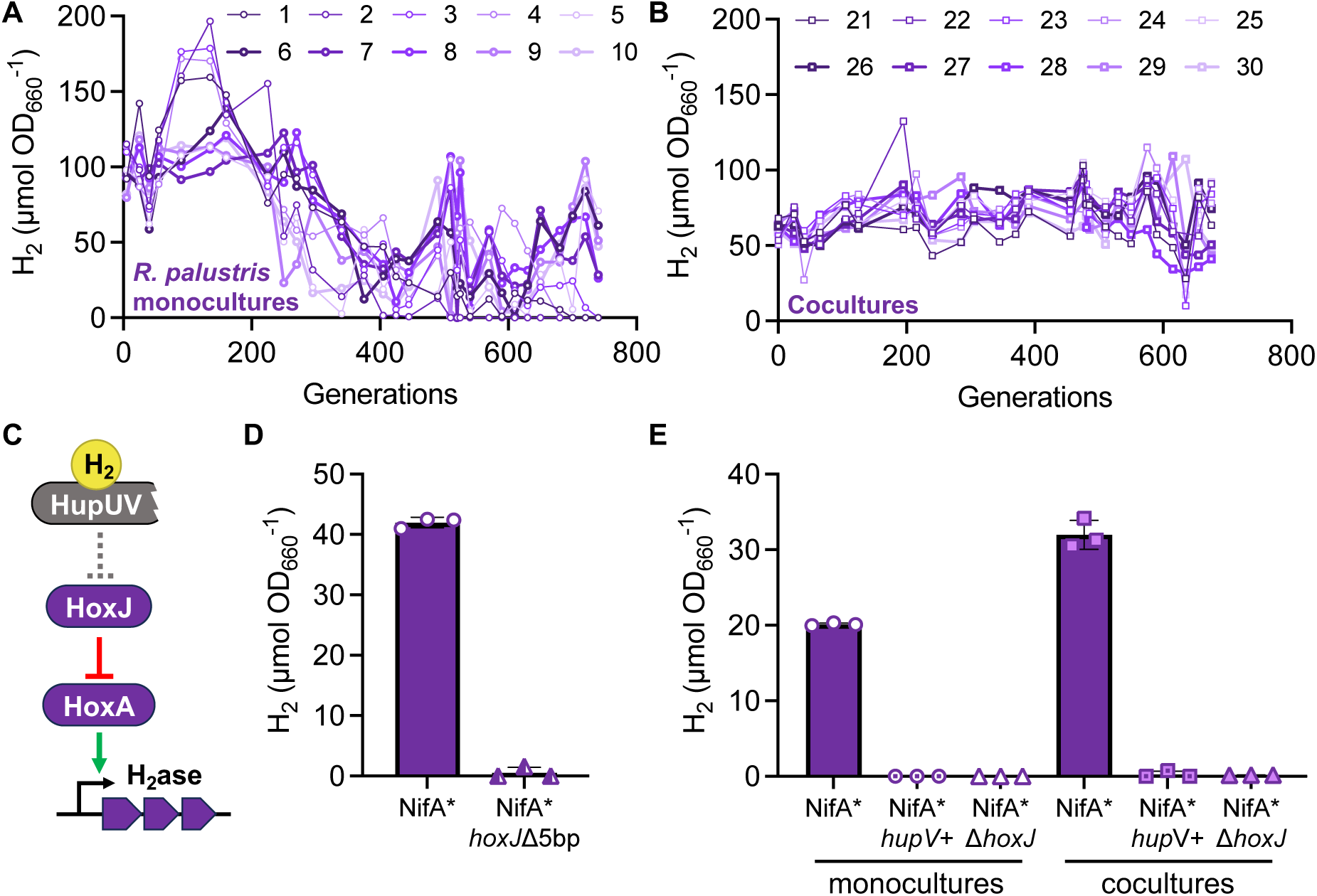
*R. palustris* monocultures evolve H_2_ utilization. H_2_ was quantified in 7-day-old cultures at selected time points for *R. palustris* monoculture lines 1-10 **(A)** and coculture lines 21-30 **(B)**. **(C)** *R. palustris* NifA* (CGA676) contains a *hupV* mutation that prevents the H_2_ sensor HupUV from repressing HoxJ. HoxJ then constitutively represses the transcriptional activator of hydrogenase (H_2_ase) genes, HoxA. **(D)** Stationary phase H_2_ in the NifA* ancestor without and with a 5-bp *hoxJ* deletion commonly observed in evolved monocultures. Cultures were grown in the same MDC medium used for monoculture serial transfers plus 1 µM NiSO_4_. **(E)** Stationary phase H_2_ levels in the NifA* ancestor without or with *hupV* repaired (*hupV*+) or *hoxJ* deleted. Monocultures were grown in MDC with 20 mM acetate. Cocultures were grown in same media used for coculture serial transfers. **(D, E)** Bars, mean ± SD; n=3.

Based on the above, we hypothesized that our *hoxJ* mutants also permitted H_2_ oxidation. To test this hypothesis, we introduced a 5-bp *hoxJ* deletion, observed in 7 monoculture lines, into the NifA* ancestor. Indeed, the mutant did not accumulate H_2_, (Fig 4D). Thus, we reason that *hoxJ* mutations that were enriched in monoculture allowed *R. palustris* to use H_2_ as an electron source.

We questioned why *hoxJ* mutants were only enriched in monoculture. Previously, transposon disruption of *hoxJ* in monoculture led to a fitness benefit in stationary phase (35). The benefit likely stemmed from H_2_-mediated CO_2_ fixation after organic substrates were exhausted; strains with WT *hoxJ* could not grow after this point (35). We first asked whether there is something about coculture conditions that generally prevents H_2_ oxidation. When we repaired the *hupV* mutation (*hupV*+) or deleted *hoxJ*, no H_2_ accumulated in either monoculture or coculture, indicating that *R. palustris* hydrogenase can be activated in coculture (Fig 4E). Therefore, something must have prevented enrichment of metabolically active *hoxJ* mutants in coculture.

We considered two possibilities that could explain the lack of *hoxJ* mutant enrichment in coculture. First, consumable organic acids, which remained available at the time of transfer in cocultures, could have allowed the general *R. palustris* population to remain competitive against *hoxJ* mutants, preventing their enrichment. A second possibility is that *E. coli* outcompeted *R. palustris* for nickel, which is required for hydrogenase activity. Since our media used for serial transfers lacked a nickel supplement, nickel must have come as a contaminant in other media components, and its availability was likely low. Although the absence of H_2_ in cocultures with Δ*hoxJ* or *hupV*+ mutants indicates that *R. palustris* can compete against *E. coli* for nickel, this might only be true when the H_2_-oxidizing *R. palustris* population is large. An emergent, low frequency, H_2_-oxidizing *R. palustris* subpopulation might instead be outcompeted by *E. coli* for nickel.

### *R. palustris* siderophore gene mutations are correlated with expression level

A frequently mutated gene cluster in *R. palustris* monocultures was RPA2382-90, for siderophore synthesis (Fig 3, Fig 5A, B). We speculate that these were LOF mutations, giving rise to ‘cheaters’ that scavenged siderophores from a siderophore-producing subpopulation. However, we questioned why *R. palustris* siderophore mutants were not observed in cocultures.

**Fig 5.**
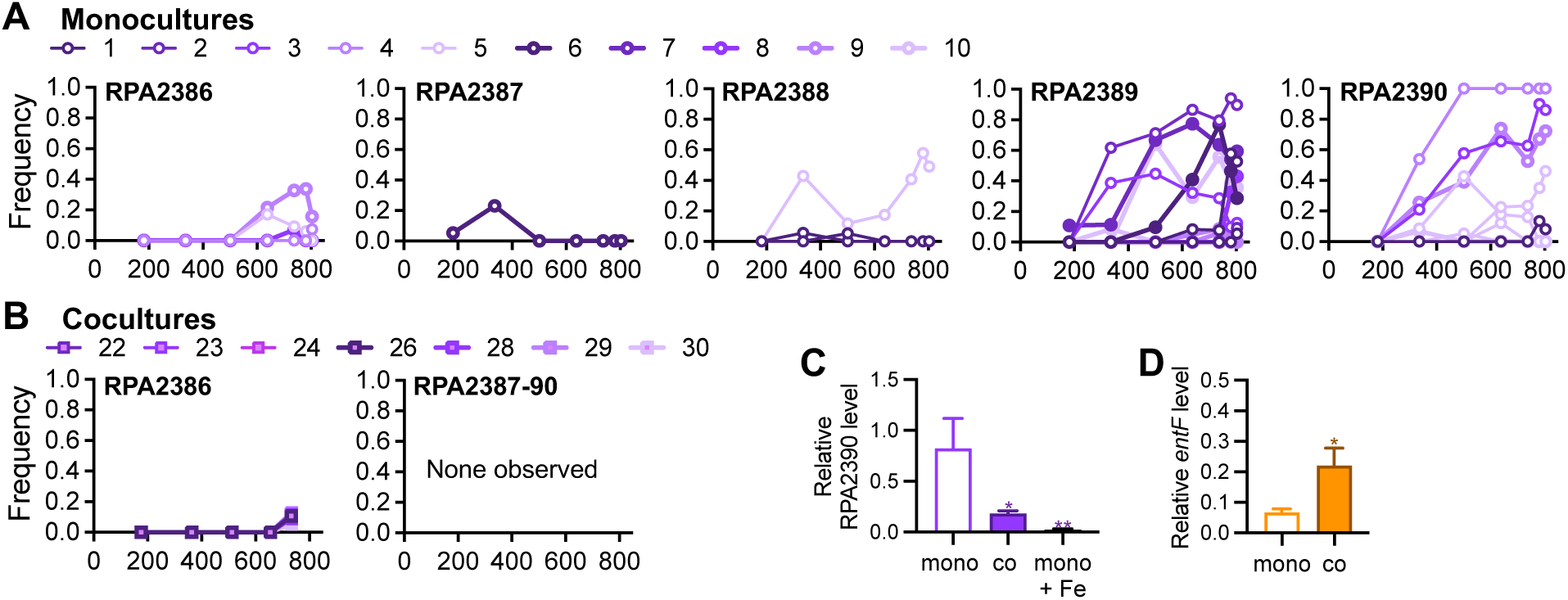
Siderophore gene loss is prevalent in *R. palustris* monocultures where gene expression is high. **A, B.** *R. palustris* siderophore gene (RPA2386-90) mutation frequencies in monoculture (**A**) and coculture lines (**B**). No mutations were observed in lines 21, 25, and 27. Repeated lines with the same symbols in a given graph indicate different mutations in the same gene. **C, D.** RT-qPCR quantification of siderophore synthesis genes in *R. palustris* (**C**; RPA2390 relative to *fixJ*) and in *E. coli* (**D**; *entF* relative to *hcaT*). Values are the mean ± SD, n = 4. Statistically significant differences from the monoculture (mono) condition for each strain were determined using an unpaired two-tail t-test; *, *p* < 0.05; **, *p* < 0.01; co, coculture; Fe, ammonium ferric citrate.

We hypothesized that siderophore gene mutations were not enriched in coculture because *E. coli* facilitated *R. palustris* iron acquisition, thus decreasing the need for *R. palustris* to synthesize siderophores. The situation would be analogous to that observed for *P. aeruginosa*, which produces siderophores only when needed, due to their cost.

High *P. aeruginosa* siderophore production in low-iron conditions led to a higher frequency of siderophore-deficient mutants compared to iron-rich conditions where siderophore production was low (12–14). In support of this hypothesis, we previously saw that *R. palustris* siderophore gene transcripts were low in coculture with *E. coli* MG1655, relative to monocultures (23). We verified that this trend was also true for *R. palustris* in coculture with *E. coli* PFM2; RT-qPCR analysis showed that the RPA2390 transcript level in coculture was 20% of that in monoculture (Fig 5C). Adding soluble iron to monocultures decreased RPA2390 expression to 3% of that observed without added iron, suggesting that the low expression in coculture was in response to higher iron availability (Fig 5C). Thus, iron must be scarce in *R. palustris* monocultures, inducing siderophore production, whereas in coculture, *E. coli* somehow increases iron availability, repressing *R. palustris* siderophore synthesis.

The mechanism of increased iron availability for *R. palustris* in coculture deserves investigation beyond the scope of the current study. However, we suspect that *R. palustris* can use *E. coli* siderophores. Others have speculated that *R. palustris* uses foreign siderophores because its genome encodes multiple siderophore transporters but only one synthesis cluster (36, 37). In support of this hypothesis, the *E. coli* siderophore *entF* transcript level was 3.3-fold higher in coculture than in monoculture (Fig. 5D), perhaps in response to iron loss to *R. palustris*.

### Other *R. palustris* mutations with unknown phenotypes

Other genes that were mutated in monoculture include a methylmalonyl-CoA mutase encoded by RPA1837, a possible flagellin encoded by RPA2297 that is outside the flagellar gene cluster, and a hypothetical protein encoded by RPA4084 (6 of 10 lines).

One of the few *R. palustris* genes that was differentially mutated in at least 5 coculture lines was RPA3297, annotated to encode a urea or short-chain amino acid transporter. Each mutation was an insertion or deletion that should cause a frame shift and thus they are all likely LOF mutations. This transporter was previously observed to have elevated protein levels in coculture compared to monoculture (23). We speculate that this elevated expression was disadvantageous, but the function of the gene product remains unknown.

Two *R. palustris* coculture lines also had a common intergenic mutation between *hbdA* and *etfA*. These genes are likely involved in fatty acid degradation and might be commonly regulated by LiuR (38). The effect of this mutation is unknown.

### Differentially enriched *E. coli* mutations mostly occurred in cocultures

The number of differentially enriched mutations in *E. coli* were oppositely skewed than for *R. palustris* (Fig 2). The same pattern held for differentially mutated *E. coli* genes and intergenic regions in at least two lineages per condition: 6 were mutated in monoculture vs 19 in coculture (Fig 6). Similar observations of cocultures leading to more mutations were made by another group who paired *E. coli* auxotrophs in reciprocal cross-feeding cocultures versus monocultures (39).

**Fig 6.**
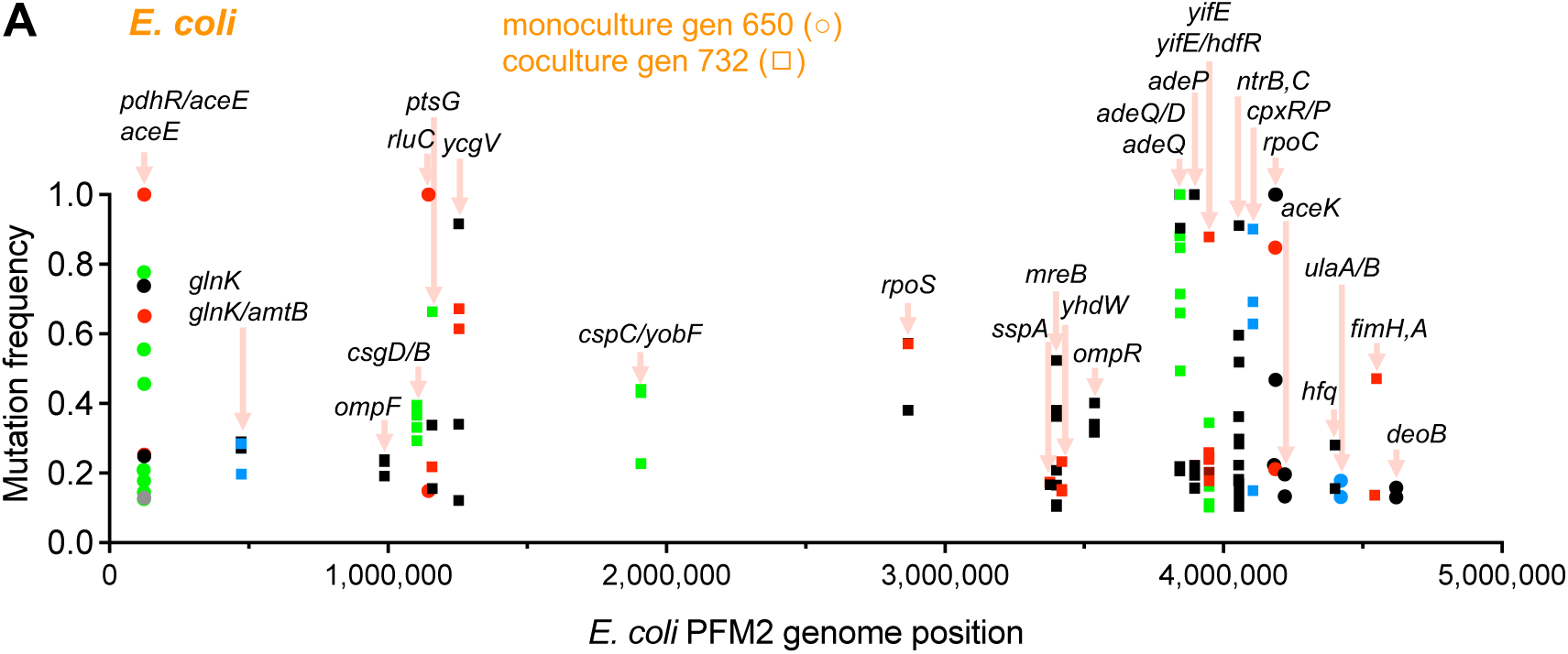
Mutations enriched in *E. coli* PFM2 in long-term monocultures and cocultures. Each point represents a mutation in a single lineage. Mutations are shown if they are not a synonymous mutation and were observed: (i) exclude synonymous mutations, (ii) no exact mutation in the opposing monoculture or coculture condition, (iii) observed in the last two sequenced timepoints, (*Ec* monoculture, gen 500 and 650; *Ec* coculture, gen 655 and 732), (iv) with a frequency >0.1 in the last timepoint, and (v) the same gene or intergenic region was mutated in at least two lines for a given condition. Generation: gen.

### *E. coli* mutations in coculture reflect adaptation to nitrogen starvation

In coculture, the *E. coli* growth rate is limited to ∼20% of maximum due to the rate at which it receives NH_4_^+^ from *R. palustris* (21, 22). As in previous long-term cocultures pairing a different *R. palustris* NifA* strain with *E. coli* MG1655 (22), several *E. coli* mutations in coculture suggested adaptation to this low-nitrogen condition. For example, mutations were commonly observed in *glnGL,* encoding the two-component system NtrBC that controls nitrogen scavenging. Previously, we demonstrated that NtrBC is critical for *E. coli* survival in the low-nitrogen coculture environment (22, 23). Other mutations in *glnK* and *amtB*, which respectively encode a nitrogen-responsive regulatory protein and an NH_4_^+^ transporter, likely also help with nitrogen acquisition in coculture (Fig 6); we also previously demonstrated the importance of *E. coli* AmtB in coculture (24).

### *E. coli* purine transporter mutants are enriched in coculture

We designed our cocultures to be based on the reciprocal exchange of fermentation products and NH_4_^+^ (21). We later discovered that *R. palustris* also externalizes adenine that can be consumed by *E. coli* (26, 27). Herein, *E. coli* exhibited several mutations in coculture that could be in response to adenine availability. For example, mutations were enriched upstream of *adeD (yicP)* encoding adenine deaminase (Fig 6). Insertion elements upstream of *adeD* in the adenine permease gene *adeQ* (yicO) are known to activate *adeD* expression (40). Mutations were also enriched in the adenine permease gene, *adeP* (*purP*) (Fig 6). The *adeP* mutations were accompanied by a multi-gene amplification that created up to 40 copies of *adeP* and neighboring genes (Fig 7A, B). We revisited sequencing results from our previous long-term cocultures featuring *E. coli* MG1655 (22) and found similar mutations upstream of *adeD* and amplification of the same region (Fig 7C). Evolved *E. coli* monocultures did not show this amplification but had up to 8-fold amplification in another region, with no obvious connection to adenine (Fig S1).

**Fig 7.**
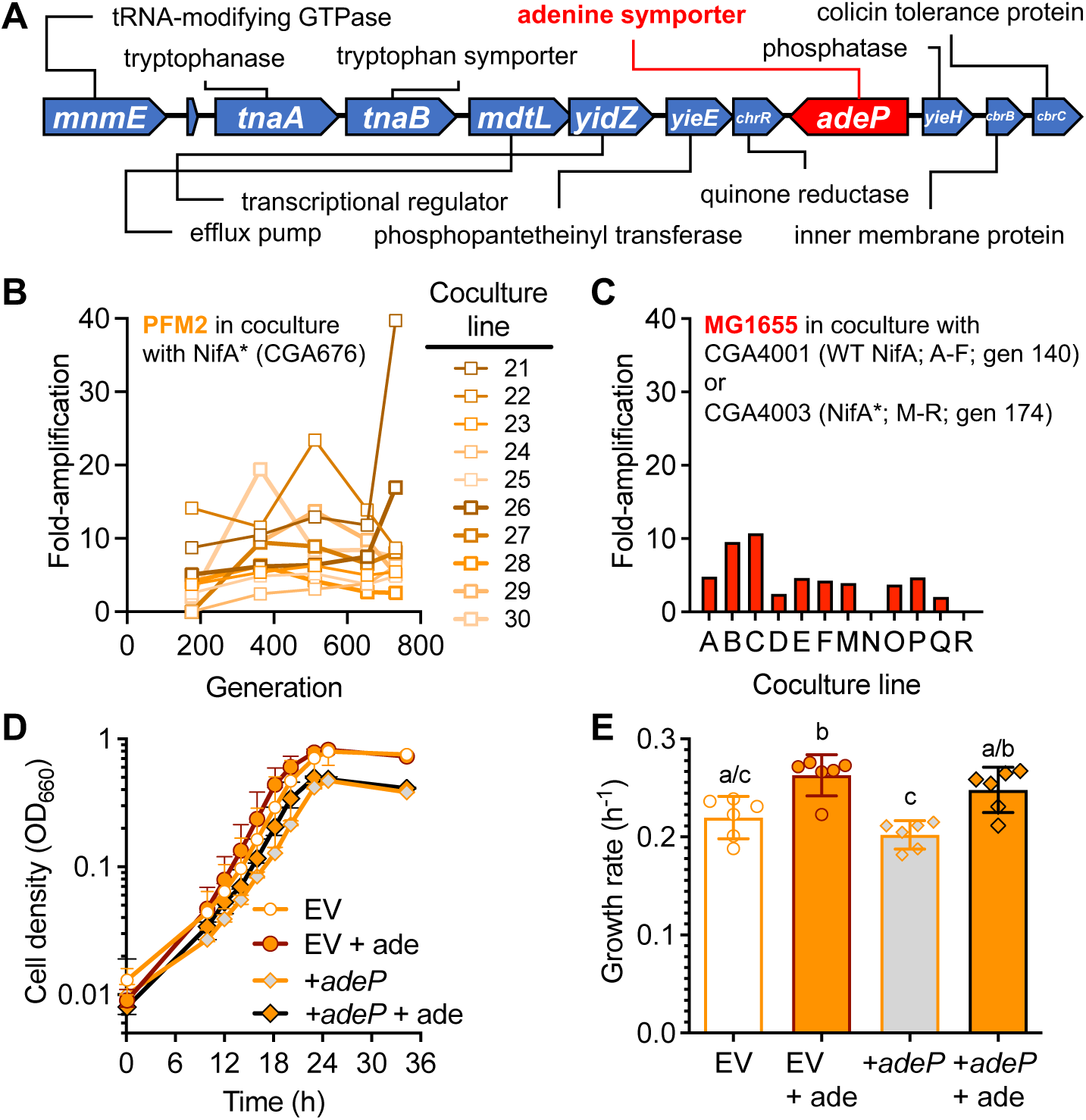
Amplification of *adeP* alone does not benefit PFM2 in the presence of adenine. **A.** Region of amplified *E. coli* genes observed in MG1655 and PFM2 cocultures but not in monocultures. The boundaries of the amplification differed between lineages but always contained these genes. **B, C.** Level of amplification observed in MG1655 in coculture (**B**) and in PFM2 in coculture (**C**). **D.** Effect of over-expressing *adeP* on PFM2 growth in monocultures with and without 35 μM adenine (ade). Over-expression was achieved by expressing *adeP* from the plasmid pCA24N with 30 μM IPTG. Data points are mean ± 95% C.I.; n=3. EV, empty vector. **E.** Growth rates from the same conditions as panel **D**; Bars, mean ± 95% C.I.; n=6, combining two separate experiments. Different letters indicate statistical differences (*p* <0.05) from a one-way ANOVA with Tukey’s multiple comparisons test.

We hypothesized that these mutations could lead to increased fitness in the presence of adenine. We attempted to mimic a high *adeP* copy number via IPTG-induced *adeP* expression from a plasmid. However, over-expressing *adeP* in PFM2 did not improve growth in the presence of adenine over an empty vector control (Fig 7D, E). It is possible that *adeP* amplification is only advantageous in the context of the surrounding amplified region or other enriched mutations.

To address an adenine-related fitness benefit more generally, we compared evolved *E. coli* growth with and without adenine. Adenine benefited several coculture-evolved PFM2 isolates, suggesting that some mutations were adaptive to adenine availability (Fig 8). Although some monoculture isolates also responded positively to adenine, including two possible auxotrophs, the trend was less pronounced; when isolates were treated as replicates, the change in cell density with adenine versus without was significantly higher for coculture-evolved PFM2 isolates but not for monoculture evolved isolates (paired one-tail t-test, *p* = 0.002 versus 0.14, respectively).

**Fig 8.**
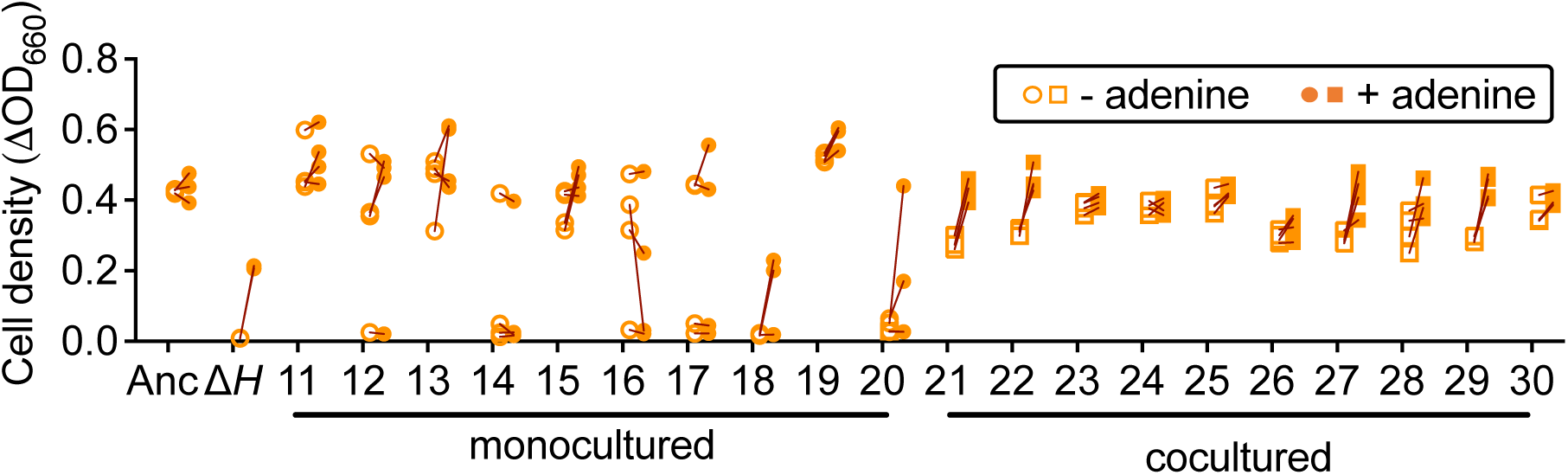
*E. coli* adenine auxotrophs are not prevalent in long-term cocultures. Randomly chosen evolved *E. coli* isolates (generation 650; n=4) were grown overnight in monoculture conditions with and without 50 μM adenine alongside the ancestral strain (Anc; PFM2) and a *ΔpurH* (*Δ*H) mutant (n=3). Each point represents a single measurement for a single isolate; replicate measurements were not made for any isolate. Each line connects measurements for the same isolate grown with and without adenine to facilitate comparisons.

### Strain-specific adenine toxicity likely influences *E. coli* evolution

Adenine availability in coculture might also explain differential enrichment of mutations in previous cocultures featuring *E. coli* MG1655 (22). MG1655 is inhibited by adenine due to an *rph* mutation that disrupts *pyrE* regulation (41), leading to suboptimal precursor distribution between purine and pyrimidine synthesis in the presence of adenine (42).

The *rph* mutation is repaired in *E. coli* PFM2 (29, 30). Through serial transfers, PFM2 mutations in *rph* and purine and pyrimidine synthesis genes were rare in monoculture and coculture (Fig 9A). In contrast, MG1655 populations (only evolved in coculture) contained a high frequency of *rph* mutations and a low frequency of purine synthesis gene (*pur*) mutations (Fig 9B). We hypothesized that MG1655 *rph* and purine biosynthesis mutations alleviated adenine toxicity. Indeed, repairing the ancestral *rph* mutation or deleting *purH*, rendering MG1655 a purine auxotroph, alleviated alanine toxicity (Fig 9C).

**Fig 9.**
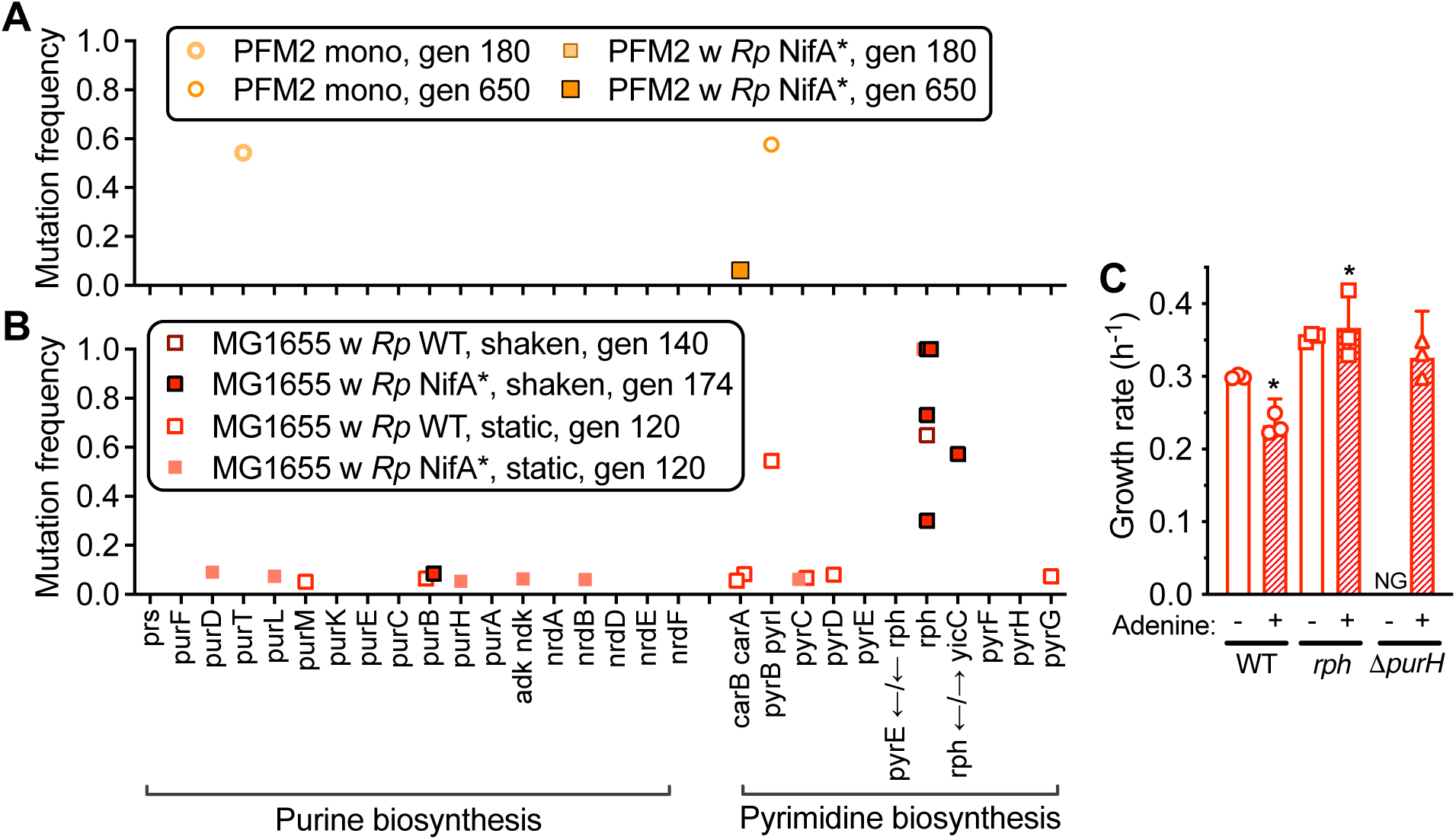
Mutations in genes for nucleotide synthesis are enriched in *E. coli* MG1655 but not PFM2. **(A,B)** Each point represents a mutation frequency in a given gene for a given line of *E. coli* PFM2 **(A)** or MG1655 **(B)**. **(C).** *E. coli* MG1655 adenine sensitivity can be relieved by repairing *rph* or deleting *purH*. Strains were grown in *E. coli* monoculture conditions without (-) or with (+) 35 μM adenine. **\***Statistical differences (*p* <0.05) compared to the wild-type (WT) conditions without adenine using a one-way ANOVA. NG, no growth.

### Coculture adenine availability did not enrich for adenine auxotrophs

Given that *de novo* purine synthesis is energetically costly, we expected to observe *E. coli* purine synthesis LOF mutations in cocultures where adenine is available from *R. palustris*; adenine is amply available in long-term cocultures (43). However, no purine biosynthesis gene mutations were observed in cocultures (Fig 9A), nor were adenine auxotrophs observed among 40 coculture isolates screened (Fig 8); only four monoculture isolates in lines 18 and 20 were possibly adenine auxotrophs (Fig 8).

The BQH predicts that cross-feeding will lead to LOF mutations only when the benefit of cross-feeding outweighs the cost of maintaining the gene (2). To directly test whether adenine auxotrophs would have an advantage in coculture, we compared PFM2 growth versus that of a *ΔpurH* mutant in a minimal medium with and without adenine. The *ΔpurH* mutant required adenine for growth, but despite the availability of adenine, the *ΔpurH* mutant did not grow faster than the PFM2 parent (Fig 10A, B). We also directly compared the fitness of PFM2 versus a *ΔpurH* mutant in coculture with *R. palustris* NifA* (CGA676). We used a range of initial frequencies so that we could simultaneously test for possible coexistence by mutual invasion, where an equilibrium frequency can be extrapolated from the x-intercept by linear regression analysis (11, 44–46). Although there was poor linear correlation in the invasion-from-rare assay, the *ΔpurH* mutant tended to decrease in frequency relative to parent in cocultures, where adenine is available (Fig 10C). Thus, our results suggest that a *ΔpurH* mutant does not have a competitive advantage over PFM2 in the presence of adenine, thus explaining why PFM2 adenine auxotrophs were not enriched in cocultures.

**Fig 10.**
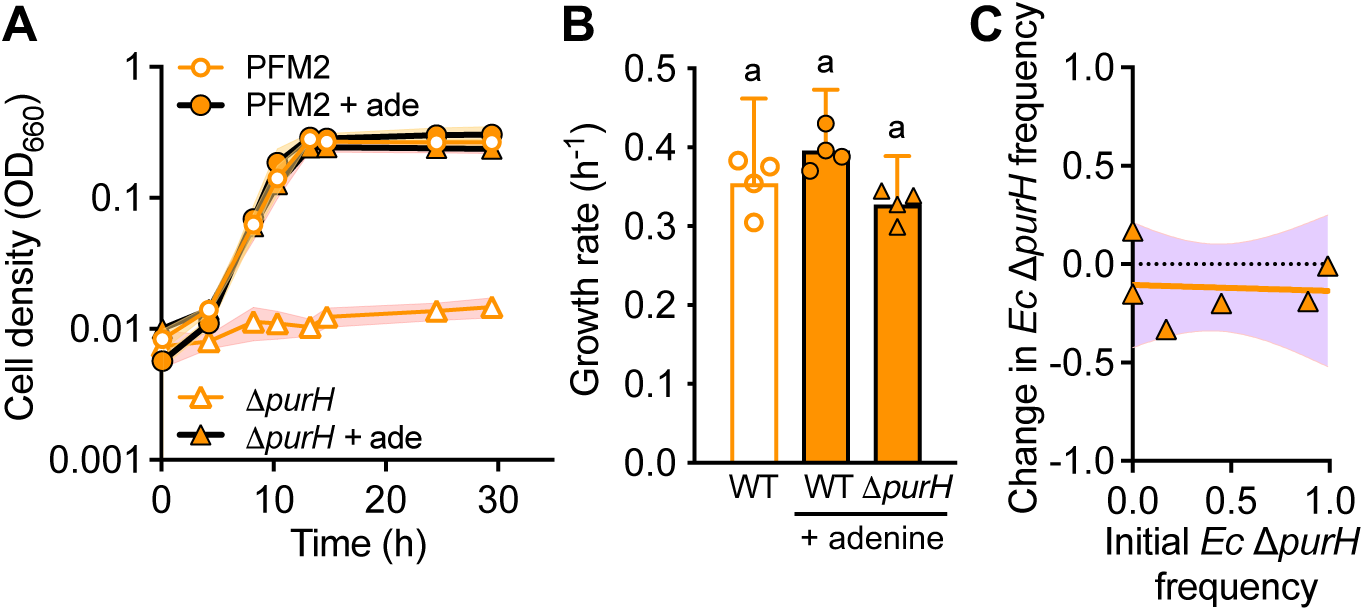
Adenine availability does not provide a fitness advantage to an engineered *E. coli* PFM2 purine auxotroph. **A.** Growth of WT *E. coli* PFM2 and its *ΔpurH* mutant in monoculture with and without 35 μM adenine (ade). Points, mean ± SD as shading; n=3. **B.** Monoculture growth rates for WT PFM2 vs a corresponding *ΔpurH* mutant with and without 35 μM adenine. ‘a,’ indicates statistically similar values (*p* > 0.01) as determined using a one-way ANOVA with Tukey’s multiple comparisons test. Bars, mean ± SD; n=3. **C.** Invasion-from-rare assay competing a *ΔpurH* mutant against its PFM2 parent in coculture with *R. palustris* CGA676 under N_2_-fixing conditions where *R. palustris* excretes NH_4_^+^ and adenine. The orange line is the best fit from a linear regression analysis with 95% CI shaded in purple. Change in frequency = (*E. coli* Δ*purH* / (*E. coli* WT + *E. coli* Δ*purH*))_final_ – (*E. coli* Δ*purH* / (*E. coli* WT + *E. coli* Δ*purH*))_initial_. **B, C.** Each data point represents a measurement from a single biological replicate.

### Adenine represses *E. coli* purine synthesis gene expression

One reason why PFM2 purine auxotrophs were not enriched in coculture could be if adenine availability repressed gene expression, thereby decreasing gene cost. This would be analogous to the situation described above for *R. palustris* siderophore genes. In support of this hypothesis, we previously saw that *E. coli* MG1655 had lower *pur* transcript levels in coculture (23). We verified that the same is true for PFM2 by RT-qPCR quantification of *purH* transcript; coculture PFM2 *purH* levels were 20% of those in monoculture (Fig 11A). Adding adenine to monocultures resulted in a *purH* transcript level that was 12% of that observed without adenine, suggesting that the low expression in coculture was due to adenine availability (Fig 11A).

**Fig 11.**
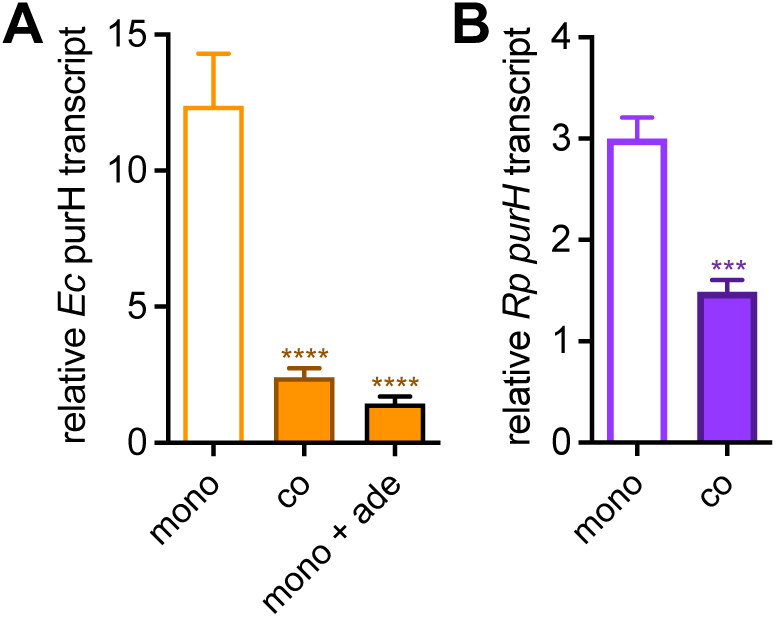
Purine biosynthesis genes are down-regulated in coculture. RT-qPCR quantification of *purH* transcript levels in *E. coli* PFM2 (**A**; relative to *hcaT*) and *R. palustris* CGA676 (**B**; relative to *fixJ*). 100 μM adenine (+ ade) was used to ensure that it was not used up before harvesting RNA. Bars, mean ± SD, n = 3-4. Statistically significant differences from the monoculture condition for each strain were determined using an unpaired two-tail t-test; ***, *p* < 0.001; ****, *p* < 0.0001.

*R. palustris* also had less *purH* transcript in coculture, 50% of that in monoculture (Fig 11B). It is unclear how *E. coli* would influence *R. palustris purH* transcript levels. Regardless, it is curious that adenine availability combined with *purH* expression in monoculture did not lead to *R. palustris pur* LOF mutations (Table S1) nor any purine auxotrophs (Fig 12). We speculate that *R. palustris* lacks adenine uptake mechanisms, which would also help explain extracellular adenine accumulation (26, 43); poor uptake mechanisms can be an important factor in extracellular metabolite accumulation (47–49).

**Fig 12.**
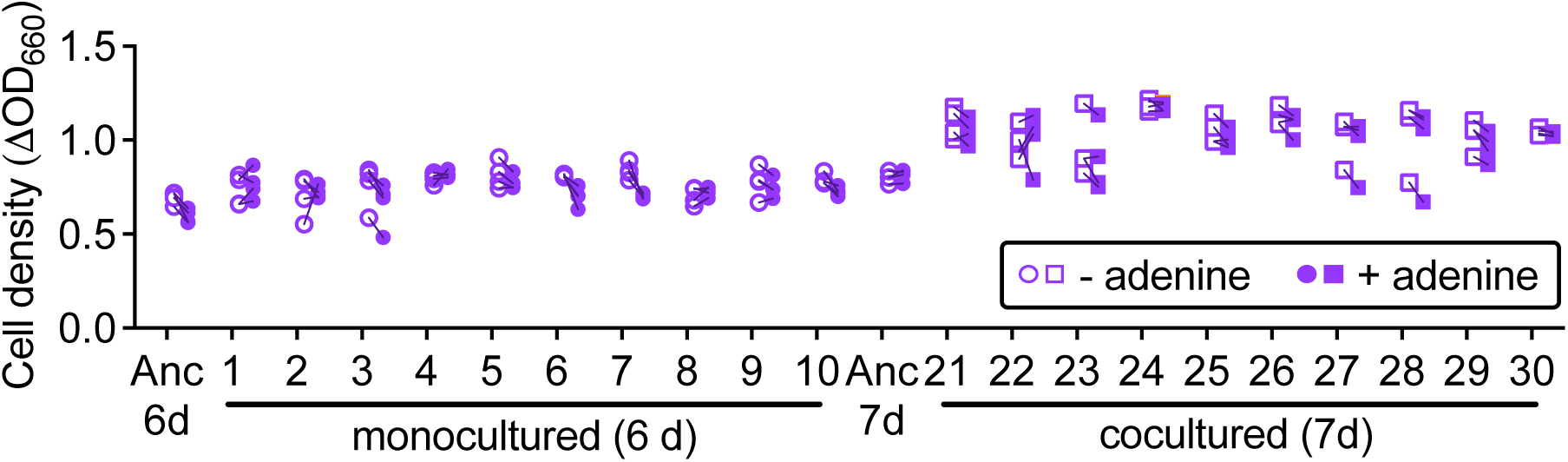
*R. palustris* auxotrophs are not prevalent in long-term cocultures. Randomly chosen evolved-*R. palustris* isolates (generation 650) were grown overnight in monoculture conditions with and without 50 μM adenine alongside the ancestral strain (Anc; CGA676) (n=4). Monoculture isolates were grown for 6 days and coculture isolates were grown for 7 days due to inclement weather, each with a corresponding ancestral control. Each point represents a single measurement for a single isolate; replicate measurements were not made for any isolate. Each line connects measurements for the same isolate grown with and without adenine to facilitate comparisons.

### *E. coli* monocultures enrich for auxotrophs

Whereas cocultures did not enrich for *E. coli* auxotrophs, several evolved monoculture PFM2 isolates could not grow in a minimal medium (Fig 8). Enrichment of auxotrophs in monoculture could suggest the emergence of cross-feeding between *E. coli* subpopulations that were not available in coculture, perhaps due to competition from an *R. palustris* population that is an order-of-magnitude larger (21, 22). Although we did not investigate these possible *E. coli* monoculture cross-feeding relationships further, we speculate that a source of the auxotrophies could be mutations in *rpoC*, encoding the RNA polymerase β’-subunit. RpoC mutations are known to result in polyauxotrophs (50).

Curiously, we did not see a similar emergence of *R. palustris* auxotrophs (Fig 12). *E. coli* and other diverse bacteria are known to support, or quickly evolve to support, auxotrophic strains via cross-feeding (51–55). It is possible that *R. palustris* cannot support diverse auxotrophs beyond those that require purines (26).

### Other *E. coli* mutations with unknown phenotypes

Other differentially enriched *E. coli* mutations include those in *aceE* that evolved in monocultures (Fig 6). *aceE* encodes a pyruvate dehydrogenase subunit, which is important for respiration but is generally not involved in *E. coli* fermentative metabolism (56). It is possible that expression of *aceE* is sub-optimally high under the high growth rates supported in monoculture conditions (23). Other mutations enriched in monoculture occurred in genes for post-translational regulation of the TCA cycle enzyme isocitrate dehydrogenase (*aceK*), ribosome modification (*rluC*), a vitamin C transporter (*ulaA/B*), and a deoxyribose salvage enzyme (*deoB*).

Several genes mutated in coculture might impact biofilm formation and acid sensitivity. We speculate that these mutations could help *E. coli* associate with *R. palustris* and/or *E. coli* cross-feeding partners and potentially respond to fermentative acid stress. For example, *csgD/B* is involved in curlin-based biofilm formation (57), which is regulated in part by *cpxRP* genes that were also mutated (58). *ycgV* encodes an outer membrane protein that might be involved in biofilm formation and *fimH/A* are involved in fimbriae-mediated adhesion (57). Mutations in *yifE* (*maoP*)/*hdfR*, might also affect biofilm formation. MaoP was noted to affect biofilm formation in addition to its role in chromosome positioning and segregation (59). HdfR might have a regulatory link to the transition between motile and sessile lifestyles (60).

HdfR might also be linked to acid resistance by regulating amino acid decarboxylation genes encoded by *gltBD* (*61*). Mutations in *hdfR* were also observed in cocultures with MG1655 (22). Mutations were also observed in *cspC*/*yobF*, which encode for proteins that respond to acid and other stressors (62). Some of the genes associated with the amplified region in cocultures might also be linked to acid resistance such as *mnmE* (63) and *yidZ* (64).

### Concluding remarks

Long-term serial transfers of *R. palustris* and *E. coli* monocultures and cocultures differentially enriched for mutations. More differentially enriched mutations were observed in *R. palustris* monocultures compared to cocultures, and vice-versa for *E. coli*. Our results suggest that the extent to which cross-feeding influences the evolutionary landscape depends on the organisms and conditions.

A generalization from our observations is that monocultures enriched for dependencies more than cocultures. Against expectations for cross-feeding to promote LOF mutations in a recipient related to synthesis of a cross-fed metabolite, *R. palustris* siderophore mutants and unknown *E. coli* auxotrophs were only enriched in monoculture. We reason that adenine and siderophore LOF mutants were not observed in coculture because cross-feeding generated metabolite levels that were sufficient to repress gene expression, thereby lowering gene cost; when gene cost is minimized, selective pressure for LOF mutations is also minimized, thus promoting gene retention. Our results suggest that the effects of cross-feeding on gene loss versus retention can be more nuanced than is often assumed.

## MATERIALS AND METHODS

### Bacterial strains

1. *R. palustris* CGA676 is derived from CGA0092 (65) and carries a *nifA*\* mutation that causes NH_4_^+^ excretion under N_2_-fixing conditions (21, 28). *R. palustris* mutants were made by homologous recombination using constructs in the suicide vector pJQ200SK (66). All primers are in Table S6. pJQhoxJΔ5bp was made by amplifying 1.4 kb from an evolved isolate from line A2 using primers JLM32 and JLM33, digesting with BamHI and XhoI sites, and then ligating into pJQ200SK that had been digested with the same enzymes. The ligation reaction was used to transform NEB10β for plasmid verification by PCR and Sanger sequencing. pJQhoxJΔ5bp was then used to transform *E. coli* S17-1 by electroporation. In each case, transformed *E. coli* was recovered on LB agar with gentamycin (30 μg ml^-1^). *E. coli* S17-1 with pJQ-ΔhoxJ and pJQ-hupV were gifts from C. S. Harwood (67).
2. *R. palustris nifA* hoxJ*Δ5bp (CGA4065), *nifA\** Δ*hoxJ* (CGA4050) and *nifA*\* *hupV*+ (CGA4062) were then made by introducing pJQhoxJΔ5bp, pJQ-ΔhoxJ, or pJQ-hupV, respectively, into CGA676 by conjugation with an *E. coli* S17-1 donor as described (67). Single recombinants were recovered on photosynthetic medium (PM) agar (68, 69) with 10 mM succinate and 100 μg ml^-1^ gentamycin. Double recombinants were recovered by counterselection on PM succinate plates with 10% sucrose, screened for gentamycin sensitivity, and then verified by PCR amplification and/or Sanger sequencing.
3. *E. coli* PFM2 was a gift from P. Foster (29, 30). The Δ*purH* mutant was made via lambda Red recombination (70) using constructs amplified from the Δ*purH*::km^R^ KEIO mutant JW3970 (71) using primers YCC29 and YCC30. FLP-mediated excision was used to remove the km resistance cassette (70). All mutants were verified by PCR amplification and Sanger sequencing.

### Growth conditions

Colonies were recovered from 25% glycerol −80 °C frozen stocks by streaking to lysogeny agar (*E. coli*) or PM agar with 10 mM succinate. Single colonies were then used to inoculate starter cultures. Anoxic media in test tubes were prepared by bubbling N_2_ through 10 ml of media in 27-ml anaerobic test tubes, then sealing with rubber stoppers and aluminum crimps prior to autoclaving. Monocultures and cocultures were grown horizontally at 30°C with light from a 45 W halogen bulb (430 lumens) and shaking at 150 rpm in minimal M9-derived coculture medium (MDC) (21) with either (i) *E. coli* monocultures: 10 mM glucose, 10 mM NH_4_Cl, and cation solution (100X stock: 100 mM MgSO_4_ and 10 mM CaCl_2_); *R. palustris* monocultures: 20 mM sodium acetate and 10 mM NH_4_Cl; or cocultures: 25 mM glucose and cation solution. Cultures with plasmid-carrying strains were also supplemented with 100 μg/ml gentamycin or 25 μg/ml chloramphenicol as appropriate. Starter cultures were inoculated with single colonies. *R. palustris* starter cultures were grown in MDC with 20 mM acetate and 10 mM NH_4_Cl. *E. coli* starter cultures were grown aerobically in lysogeny broth, with 30 μg/ml kanamycin (km) when appropriate. *E. coli* starter cultures were washed twice in 1 ml MDC prior to inoculating test cultures or bioassays. Cocultures were inoculated with 0.1 ml each of *R. palustris* and *E. coli* to an initial optical density (OD_660_) of ∼0.003 each.

Cultures in 96-well plates were treated similarly except that oxic solutions were used to prepare 0.2 ml volumes in each well. Anoxic conditions were achieved by sealing plates inside a BD GasPak EZ large incubation container with anaerobic sachets.

### Experimental evolution

Founder monocultures of *E. coli* PFM2 and *R. palustris* NifA* CGA676 were each grown from a single colony in anoxic MDC with either 10 mM glucose, cation solution, and 3 mM NH_4_Cl for PFM2 or 20 mM sodium acetate for CGA676. A single founder monoculture was then used to inoculate 10 monocultures and 10 cocultures. All cultures were grown horizontally without shaking at 30°C with light in MDC. PFM2 monocultures were supplemented with 10 mM glucose, cation solution, 25 mM NaCl, and 2.3 mM NH_4_Cl. These cultures were started later after finding that 25 mM glucose led to periodic extinction due to prolonged exposure to acidic fermentation products. As a result, *E. coli* monocultures were transferred less times than for other conditions. No data from monocultures with 25 mM glucose were presented herein. CGA676 monocultures were supplemented with 25 mM glucose, cation solution, 8 mM disodium succinate, 7.3 mM sodium acetate, 0.25 mM sodium formate, 1.4 mM sodium lactate, and 6.3 mM ethanol. Cocultures were supplemented with 25 mM glucose, cation solution, and 25 mM NaCl. Acidification was not a problem despite using 25 mM glucose because *R. palustris* consumes enough organic acids to dampen the pH drop (21). Every 7 days, cultures were vortexed and 0.25 ml was transferred to fresh medium. About every 5 transfers, stocks were diluted 1:1 with 50% glycerol and stored at −80°C. Separately, cell pellets from 1 ml samples were frozen for gDNA extraction.

### Analytical procedures

Cell densities were measured via turbidity (OD_660_) using a Genesys 20 visible spectrophotometer (Thermo-Fisher). H_2_ was measured using a Shimadzu GC-2014 gas chromatograph with a thermal conductivity detector as described by sampling 0.1 ml of headspace using a gas-tight syringe (72). Statistical analyses were performed using Graphpad Prism (v10).

### Competition assays

Competition assays were conducted in an invasion-from-rare format to consider coexistence by the mutual invasion criterion, where each population can increase when rare (44, 46). Cocultures were started from various initial frequencies (targeting 0.01 – 0.99) of each *E. coli* strain (WT vs Δ*purH* or WT vs Δ*purH*::km^R^) for a total initial cell density of ∼10^6^ colony forming units (CFU) / ml. *R. palustris* CGA676 was inoculated to an initial density of ∼10^6^ CFU / ml. Frequencies were determined upon inoculation and after 5 days. WT and Δ*purH* mutants were distinguished by counting CFUs on M9 agar with cations and 25 mM glucose and on LB agar and then determining Δ*purH* populations from the difference. When Δ*purH*::km^R^ mutants were used LB agar included km to allow for direct determination of population size. Change in frequency = (*E. coli* Δ*purH* / (*E. coli* WT + *E. coli* Δ*purH*))_final_ – (*E. coli* Δ*purH* / (*E. coli* WT + *E. coli* Δ*purH*))_initial_ (45).

### Genome sequencing and mutation analysis

gDNA was purified from cells using a Qiagen DNeasy Blood and Tissue kit following the manufacturer’s instructions. Lysis was facilitated after resuspension by adding proteinase K (50 μg/ml final), and incubating at 56°C for 10 min. RNaseA (4 μl, Promega) was then added and the lysate was incubated for 2 min before proceeding.

DNA fragment libraries were made using a NextFlex Bioo Rapid DNA kit and libraries were sequenced using Illumina NextSeq 500 150×150 paired-end runs by the IU Center for Genomics and Bioinformatics. Paired-end reads were pre-processed for quality with cutadapt 3.4 (73) with the following options: -a AGATCGGAAGAGC -A AGATCGGAAGAGC; -q 15,10; -u 6. Mutations were called using breseq v. 0.32.0 on polymorphism mode (74). *E. coli* monoculture population sequences were mapped to the MG1655 genome (accession NC_000913); *R. palustris* monoculture population sequences were mapped to a concatenated reference genome consisting of the CGA009 chromosome (accession BX571963), and its plasmid pRPA (accession BX571964). Co-culture sequences were mapped to a concatenation of the *E. coli* and *R. palustris* reference genomes. Polymorphisms that co-occurred in both the monoculture and co-culture datasets were filtered and maintained as a subset to enrich for the most informative variants representing treatment differences (Table S1-4). This filtering step also removed sequence differences between the reference sequences and those of the experimental strains used. Variants were prioritized as mutations of interest if they were detectable at the final two sequencing timepoints and co-occurred across multiple populations in the same locus. All mutations can be found in Tables S1-4. Locus tag conversions can be found in Table S5.

### Reverse transcription quantitative real-time PCR (RT-qPCR)

Cultures received 100 μM adenine or ammonium ferric [iron(III)] citrate as indicated. Cultures were harvested in exponential phase at 0.6-0.8 OD_660_ except for *E. coli* monocultures (+/- adenine experiment), which were harvested at 0.3-0.4 OD_660_ to avoid adenine depletion.

Cultures were chilled on ice and pelleted by centrifugation. Lysis, RNA purification, and cDNA generation were performed exactly as described (26). Standard curves were generated using gDNA. Transcripts were quantified as described (26) using the appropriate primers (Table S6) with a Mastercycler ep *realplex* real-time PCR system (Eppendorf). Data was analyzed by *realplex* software using Noiseband. Specificities were validated by melting curves and by the presence of a single band on an agarose gel.

## ACKNOWLEDGEMENTS

This work was supported in part by US Army Research Office grants W911NF-14-1-0411 and W911NF-17-1-0159, the National Science Foundation CAREER award MCB-1749489 to JBM, and National Institutes of Health grants R35GM128674 to ABD and R35GM150625 to MB. Supercomputing resources were supported in part by Lilly Endowment, Inc., through its support for the IU Pervasive Technology Institute.

We are grateful to A. Cairo, S. Kamaran, and N. Ramli, for maintaining long-term cultures. We thank A. Drummond, J. Drummond, P. Foster, J.P. Gerdt, J. Lennon, M. Lynch, T. Romeo, and the McKinlay lab for helpful advice.

