## Supplementary material for "Reciprocal cross-feeding between bacteria can limit the emergence of metabolic dependencies": Fig S1

**CONTENTS:**

**Table S6. Primers**

**Fig S1. *E. coli* PFM2 gene amplification region in monocultures**

**Table S6. Primers.**

| Primer | Sequence (5'-3') | Description |
|---|---|---|
| JLM32 | tagtggatccgctcaccgatctcgtcgatc | Forward *ΔhoxJ*5bp (BamHI) |
| JLM33 | actgCTCGAgagtagcggtcggacgttc | Reverse *ΔhoxJ*5bp (XhoI) |
| YCC29 | gcgcaaacgttttcgttacaatgcg | 5' of Δ*purH*::*Km* in JW3970 |
| YCC30 | tgcattacccggagcaac | 3' of Δ*purH*::*Km* in JW3970 |
| YCC80 | taagggaaccgtgcatgtg | Forward qPCR primer for *Rp fixJ* |
| YCC81 | ggattcgtacagcttgacctc | Reverse qPCR primer for *Rp fixJ* |
| YCC97 | acgtcgtcgtggttcttg | Forward qPCR primer for *Rp purH* |
| YCC98 | cgaagccaccgtcgataaa | Reverse qPCR primer for *Rp purH* |
| YCC99 | gcaacacgttctgctgatg | Forward qPCR primer for *Rp RPA2390* |
| YCC100 | cattggttctcggcctatct | Reverse qPCR primer for *Rp RPA2390* |
| YCC91 | aaccgcatggcccttatt | Forward qPCR primer for *Ec entF* |
| YCC92 | gtatccagcaagccaagaaatg | Reverse qPCR primer for *Ec entF* |
| YCC93 | gtgttcgaaggctttgatgg | Forward qPCR primer for *Ec purH* |
| YCC94 | gtgaagagcagcgactatga | Reverse qPCR primer for *Ec purH* |
| YCC95 | gtcgcactttgccgtaatc | Forward qPCR primer for *Ec hcaT* |
| YCC96 | gctgatgctggtgatgattg | Reverse qPCR primer for *Ec hcaT* |

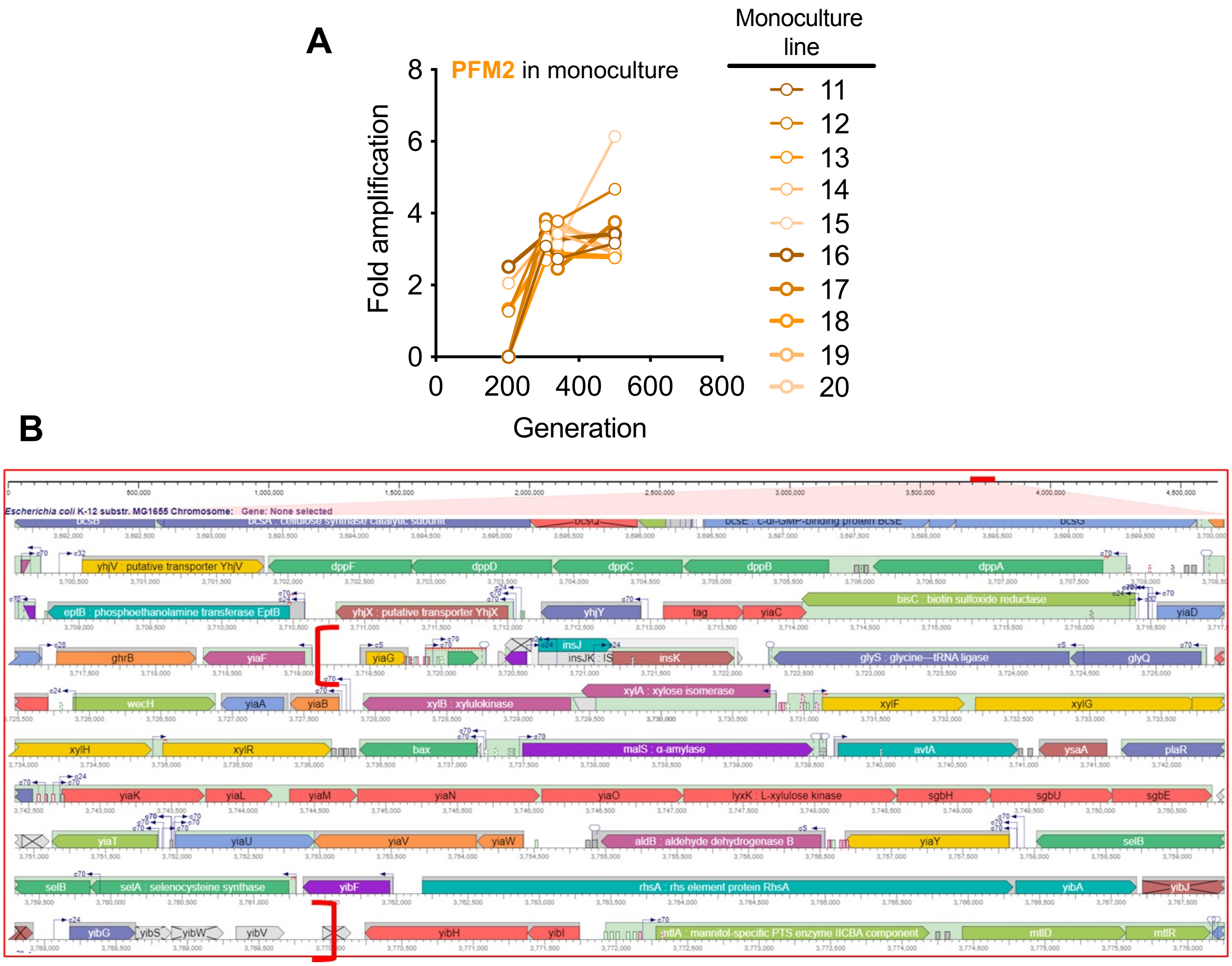


**Fig S1.** ***E. coli*** **PFM2 gene amplification region in monocultures. A.** Level of amplification observed in PFM2 monocultures. **B.** Amplified region of the chromosome based on mapping to *E. coli* MG1655. Image was generated using Metacyc.org.
